# Fixed-Target Serial Crystallography at Structural Biology Center

**DOI:** 10.1101/2022.04.06.487333

**Authors:** Darren A. Sherrell, Alex Lavens, Mateusz Wilamowski, Youngchang Kim, Ryan Chard, Krzysztof Lazarski, Gerold Rosenbaum, Rafael Vescovi, Jessica L. Johnson, Chase Akins, Changsoo Chang, Karolina Michalska, Gyorgy Babnigg, Ian Foster, Andrzej Joachimiak

**Affiliations:** Structural Biology Center, X-ray Science Division, Argonne National Laboratory, Lemont, Illinois 60439, USA; Center for Structural Genomics of Infectious Diseases, Consortium for Advanced Science and Engineering, University of Chicago, Chicago, Illinois 60667, USA; Data Science and Learning Division, Argonne National Laboratory, Lemont, Illinois 60439, USA; Biosciences Division, Argonne National Laboratory, Lemont, Illinois 60439, USA; Department of Biochemistry and Molecular Biology, University of Chicago, Chicago, Illinois 60367, USA

**Keywords:** fixed-target serial synchrotron crystallography

## Abstract

Serial synchrotron crystallography enables studies of protein structures under physiological temperature and reduced radiation damage by collection of data from thousands of crystals. The Structural Biology Center at Sector 19 of the Advanced Photon Source has implemented a fixed-target approach with a new 3D printed mesh-holder optimized for sample handling. The holder immobilizes a crystal suspension or droplet emulsion on a nylon mesh, trapping and sealing a near-monolayer of crystals in its mother liquor between two thin mylar films. Data can be rapidly collected in scan mode and analyzed in near real-time using piezoelectric linear stages assembled in an XYZ arrangement, controlled with a graphical user interface and analyzed by using a high-performance computing pipeline. Here, the system was applied to two *β*-lactamases: a class D serine *β*-lactamase from *Chitinophaga pinensis* DSM 2588 and L1 metallo-*β* -lactamase from *Stenotrophomonas maltophilia* K279a.

## 1. Introduction

Synchrotron serial crystallography (SSX) has emerged as a valuable approach for low-dose room-temperature structural biology research that also allows for the study of dynamic processes in protein crystals, such as chemical transformations. SSX followed and refined serial crystallography experiments ^1^ performed at x-ray Free-Electron Lasers (XFELs). The need arose because the powerful pulses from XFELs destroy each crystal after a single exposure, and thousands of crystals must be exposed to capture enough diffraction reflections to solve and refine the structure ^2,3^. Improvements in technology since the introduction of serial crystallography have made SSX much more accessible and routine ^4-11^. SSX has become available at many microfocus synchrotron beamlines and is being widely adopted by the structural biology community.

While the serial crystallography field has matured significantly ^1^, challenges remain ^12,13^. Sample preparation and delivery is the critical element in these experiments, and various approaches have been developed ^8,14-19^. Existing solutions for serial crystallography include: i) liquid injectors (fast and slow) ^18,20-22^, ii) fixed-target including silicon-nitride ^8^ and micro-patterned chips ^19^, iii) microfluidic traps ^17^, iv) nylon mesh ^16^, v) “chipless chips” ^14^, and vi) tape-drives ^23^. Each method has unique advantages, such as mixing and releasing compounds, delivery in a vacuum, temperature ranges, humidity-control, and delivery speed. Drawbacks can include high sample consumption, difficulty in loading, and timing considerations, as well as overall complexity that may create a barrier to use.

In addition to delivery methods and environmental options, an essential driver for serial crystallography experiments is biology and biochemistry. SSX can capture structural changes, including binding of metals, molecules, and cofactors, and recording chemical transformation, including enzymatic catalysis, in a time-dependent manner (manuscript in preparation). Structures measured at different time points can help visualize structural and chemical changes with time scales spanning sixteen orders of magnitude, from femtoseconds ^2^ to minutes ^24^. While it still takes thousands of crystals to measure each time point, the advancements in modern light sources, delivery methods, detectors, and analysis pipelines have converged to make atomic-resolution structural dynamics an exciting new investigative option.

We have recently demonstrated the benefit of the SSX approach in studies of the SARS-CoV-2 Nsp10/16 2′-O RNA methyltransferase complex. The crystal structures of Nsp10/16 with substrates (Cap-0 analog and S-adenosyl methionine) and products (Cap-1 analog and S-adenosyl-L-homocysteine) revealed the states before and after methylation, occurring within the crystals during the experiments ^11^. Our previous publication focused on a scientific case; here, we describe the approaches to better utilize light sources x-ray beams and biological samples that we have developed to meet increasing demand for reliable and user-friendly access to SSX capabilities. We report a fixed-target mesh delivery method that is simple, low-cost, accepts crystals of various sizes, and can be applied to both static and time-resolved studies. We have implemented this SSX data collection system at the Structural Biology Center (SBC) 19-ID beamline at the Advanced Photon Source (APS). Our method employs the high-precision, high-speed motorized stages introduced by Owen et al. ^25^. This report emphasizes the newly designed holders which are 3D printed, low-cost, reliable, and can be quickly loaded and reused. The sample environment is a mylar-mesh-mylar sandwich (Fig. 1), sized to enable collection of a complete diffraction data set from a single holder under standard conditions. Crystal loading requires only a steady hand and a regular pipette, the holders are attached onto the end-station goniostat by using a magnetic mount in the same fashion as a standard single crystal crystallography pin. Data collection is controlled by simple software developed at SBC and downstream image processing engages a supercomputer-based pipeline that carries out all necessary steps in the structure determination protocol – from image integration to phasing via molecular replacement – as fast as data are collected at the beamline.

**Figure 1.**
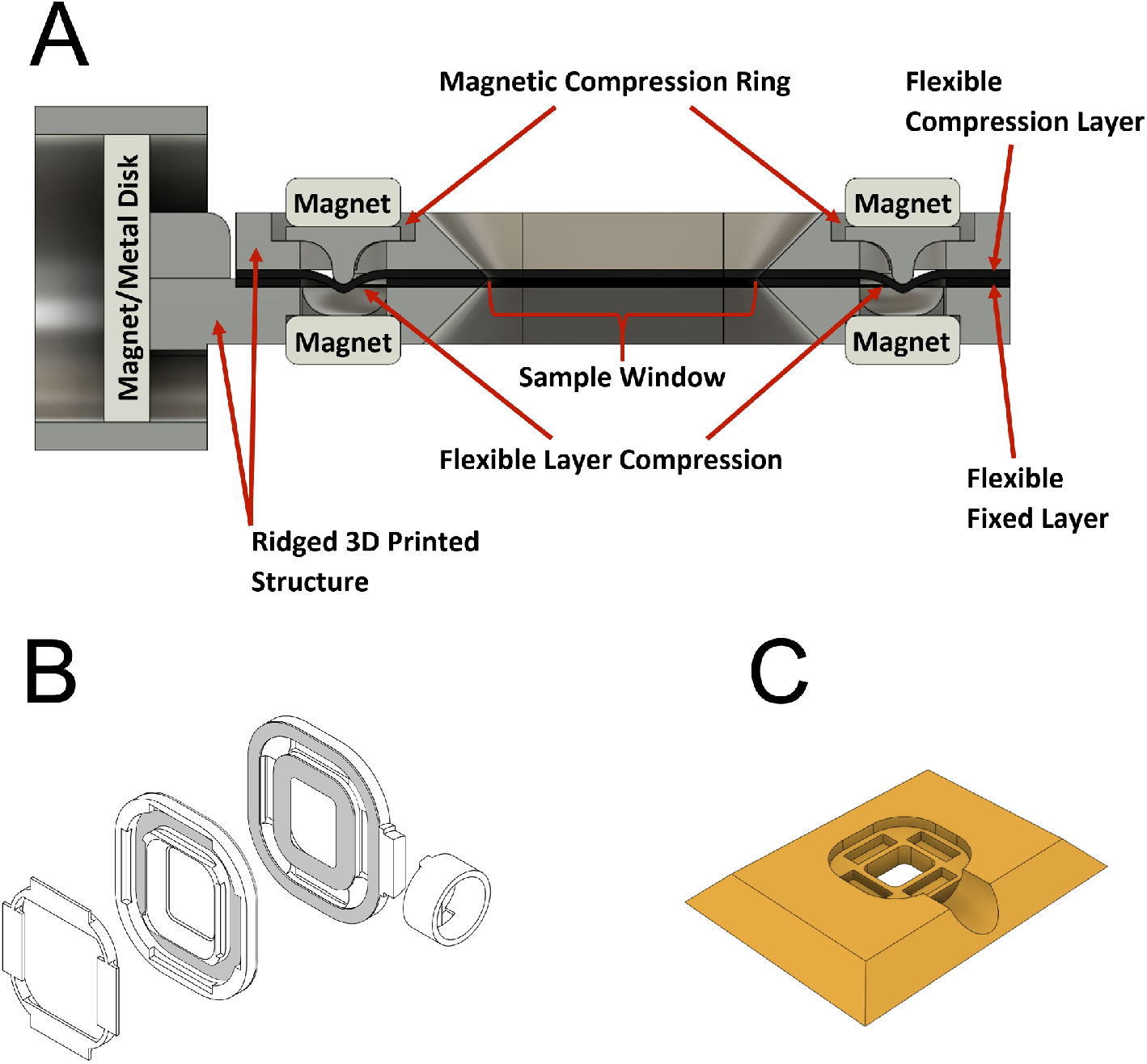
*ALEX* mesh holder (A) Cut-away view of mesh holder, (B) top and (C) isometric view of mesh holder loading plate.

We undertook the SSX approach having the general user community in mind such that serial and dynamic x-ray crystallography can become more accessible to the broader population of researchers. We demonstrate our technique’s successful use by highlighting crystal structures of two model proteins solved previously under cryo-conditions at the SBC – class D *β*-lactamase (DBL) from *Chitinophaga pinensis* DSM 2588 and L1 metallo-*β* -lactamase (MBL) from *Stenotrophomonas maltophilia* K279a. Since 2019, several successful SSX experiments have been performed at SBC, including research on COVID-19 projects ^11^.

## 2. Methods

### 2.1. Beamline set up

19-ID uses the Advanced Photon Source (APS) undulator A to generate x-rays and a Rosenbaum double-crystal monochromator design ^26^ with a constant x-ray beam height and a sagittally bent second crystal for horizontal focusing. The beamline is typically operated at energies between 6.0 and 19.5 keV. Together with the 107 mm vertical-focusing mirror, 19-ID delivers a maximum 3×10^12^ ph/s fully focused at 40 (v) x 83 (h) micrometers which are slit down further depending on sample requirements ^26^. Since the collection of the data mentioned here and to minimize the impact of SSX experiments on regular beamline operations, the developed SSX motors for fixed-target stages have been permanently integrated into the end-station, rather than moved into place as needed, to be used for crystal alignment with the x-rays during ‘normal’ single-crystal experiments (Fig. 2). This set up allows collection of thousands of diffraction still images in a semi-automated scan mode. An additional upgrade since these mentioned experiments has been the implementation of a compact refractive lens (CRL) system, enabling beam sizing down to approximately 15×5 μm ^27^.

**Figure 2.**
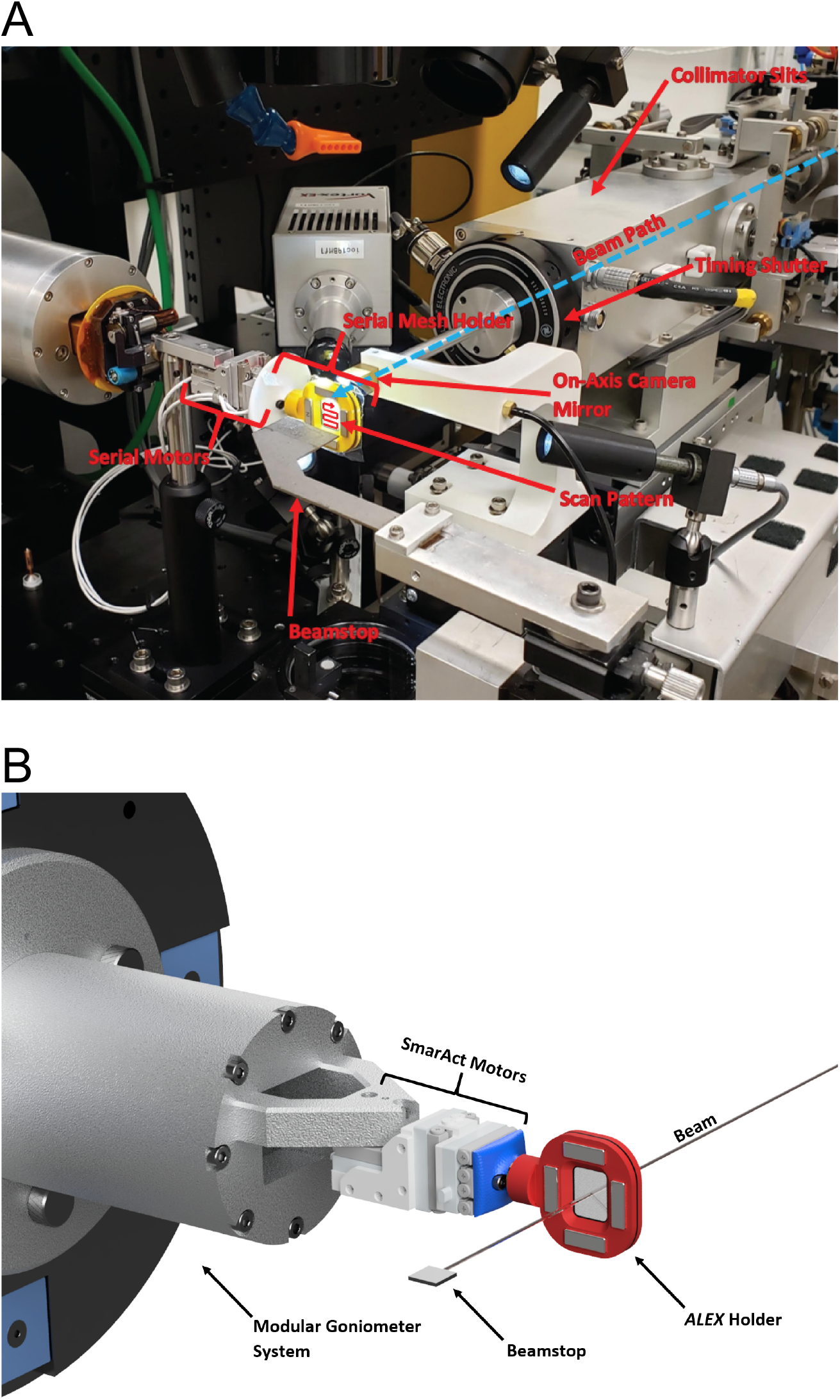
Set up of *ALEX* at 19-ID beamline (A) using modular magnetic mount prior to permanent integration onto goniostat (used in data collection reported here), (B) and schematic of permanent integration onto goniostat with upgraded configuration.

### 2.2. End station configuration

SBC uses SmarAct SLC-1720 piezoelectric linear stages assembled in an XYZ arrangement for both serial crystallography and standard cryo-crystallography sample manipulation. The XYZ configuration allows a maximum of 9 and 12 mm of travel for the X and Y directions, perpendicular to the beam, and 12 mm for Z, the x-ray beam direction. The system has linear range to perform various SSX experiments and is compact enough for single-crystal cryo-crystallography sample translation. The system movement is controlled via SmarAct’s SDC2 controller, which receives stepper motor signals from a Delta-Tau Turbo PMAC2 PCI ACC-24E2S stepper motor card. The detector is set to the external trigger mode, it receives an initial shutter signal from the Programmable Multi-Axis Controller (PMAC) and the data collection is carried out in a shutterless mode. X-ray shutter and detector image triggers are controlled using 5V Transistor-Transistor Logic (TTL) pulses from the PMAC accessory 48 I/O card. The detector can take a single or multiple images at each stopped location if a Multiple Serial Structures (MSS) ^28^ or radiation damage experiment ^29^ is desired. A long-range cryostream retractor is integrated to vacate the sample area leaving a large open volume for SSX and other experimental work.

### 2.3. SSX *sample holder and loading plate*

The SBC adopted a fixed-target approach, initially with a silicon chip ^25^ and later with a nylon mesh (this work). In the latter approach, the crystals suspension is applied on a nylon mesh and sealed between two 6 μm mylar films. A wide range of crystal sizes can be trapped and immobilized as a near-monolayer in its mother liquor between the films. The mylar-mesh-mylar sandwich is held in place and sealed by a newly designed sample holder coined Advanced Lightweight Encapsulator for Crystallography (*ALEX)* (Fig. 1A and B).

*ALEX* utilizes a novel compression seal (USPO Patent Application #16/903,601) and is made using a dual-nozzle 3D printer. The main sections of the *ALEX* holder are fabricated by 3D printing a flexible NinjaFlex (TPU 85A) layer and subsequent deposition of a ridged material (nylon, TPU-75D, etc.) structure, creating a two-layered composite material. The compression ring, pin base, and loading plate are printed from only the ridged material. The mylar-mesh-mylar sample sandwich is placed between the two composite pieces. When the two sides are brought together, the compression ring is placed and draws the two halves together, flexing the NinjaFlex layer to create an airtight seal that runs the sample area’s circumference (Fig. 1 and Supplementary Figure 1). The NinjaFlex material also provides a mildly adhesive surface which helps the mylar remain stationary. All parts of a single holder can be 3D printed in under an hour (Fig. 1).

A user-friendly loading plate was developed (Fig. 1C) for the *ALEX* holder to streamline the loading process and lower the likelihood of error and loss of sample. To load the holder, the main section of the *ALEX* holder is placed into the impression in the loading plate, making it flush with the surface. The initial layer of mylar is then placed with the nylon mesh following. The crystal suspension is then applied to the mesh by pipette and excess liquid is removed by blotting. The second mylar film is laid over the mesh/solution and the second half of the holder is placed on top, creating a holder-mylar-sample/mesh-mylar-holder structure. The magnetic compression ring is then placed, sealing and holding the entire device together, ready for data collection. Excess mylar film at the edges can be trimmed as necessary.

Currently, *ALEX* holders come with two window sizes (12.4 × 9.4 or 7 × 7 mm^2^) and can collect a sufficient number of diffraction images for a protein structure under normal conditions. The remaining parameters of the holder are physically minimized while still allowing for reliable function of the device. The window’s center is located approximately 18.5 mm from the XYZ stage mounting surface, which corresponds to the standard length of a crystallographic pin. Such characteristics, together with the implemented mounting base, allow for placement of the device on the same magnet as a standard pin, enabling installation as-is on other synchrotron beamlines. Sample area, distance from the base, and overall geometry of the holder can be adjusted to allow for use on stages with longer ranges or attachments with other devices to suit various needs.

The complete set-up was tested and commissioned at 19-ID beamline with various crystal sizes (10 – 120 μm) and morphologies (rods, needles, cubes etc.) (Fig. 3, Fig. S1).

**Figure 3.**
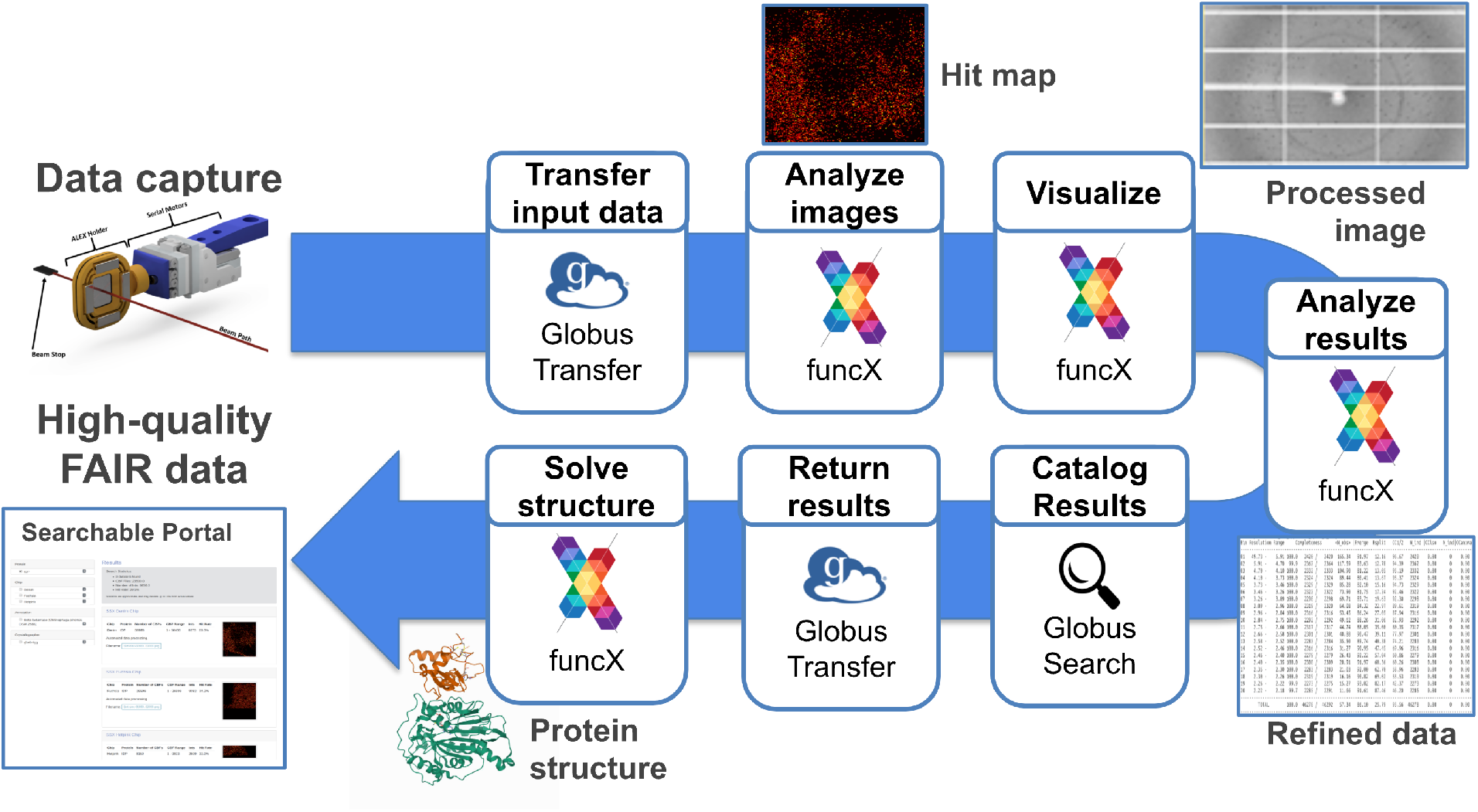
Data collection and processing pipeline. Data captured at 19ID trigger a Globus Flow to transfer images to Argonne’s Leadership Computing Facility where they are analyzed using the Theta supercomputer. Results are visualized and published to a searchable data portal for review.

## 3. Sample, purification and batch crystallization

### 3.1. Protein purification

The L1 MBL and DBL have been cloned to pMCSG7 that carries N-terminal, removal His_6_-tag, which upon cleavage leaves three artifact residues (Ser-Asn-Ala) at N-termini.

The *E. coli* BL21(DE3)-Gold strain (Stratagene) over expressing L1 MBL (residue 20-290) from *S. maltophilia K279a* was grown as described previously ^30^. Specifically, a one-liter culture of enriched LB medium was grown at 37 ^°^C (190 rpm). Bacterial cells were harvested by centrifugation at ×7000 g (Sorval evolution RC centrifuge, Thermo Scientific) and frozen. Frozen cells were thawed, resuspended in a 35 ml lysis buffer (500 mM NaCl, 5 % (v/v) glycerol, 50 mM HEPES pH 8.0, 20 mM imidazole and 10 mM β-mercaptoethanol) with a protease inhibitor cocktail tablet (Complete Ultra-EDTA-free, Sigma) per liter culture, treated with lysozyme (1 mg/ml) and sonicated (5 min total time, 130 W power output). Cells were spun at 30,000 × g at 4 ^°^C for 1 h.

The *E. coli* BL21(DE3)-Gold strain (Stratagene) overexpressing DBL from *C. pinensis* DSM 2588 (residues 21-267, accession No. ACU58405.1) were grown at 37 ^°^C and shaken at 200 rpm in enriched M9 medium until an OD @600 nm of 1.0 was reached, as described before ^31,32^. Methionine biosynthetic inhibitory amino acids (25 mg/l each of L-valine, L-isoleucine, L-leucine, L-lysine, L-threonine, L-phenylalanine) and L-selenomethionine (SeMet, Medicillin) were added and the cultures were transferred to 4 ^°^C for 1 hour before bringing them up to 18 ^°^C. The protein expression was induced with 0.5 mM isopropyl β-D-1-thiogalactopyranoside (IPTG). The cells were incubated overnight, harvested and resuspended in lysis buffer (500 mM NaCl, 5 %(v/v) glycerol, 50 mM HEPES–NaOH pH 8.0, 20 mM imidazole, 10 mM 2-mercaptoethanol).

Both proteins were purified by immobilized metal-affinity chromatography (IMAC-I) on an ÄKTAxpress system (cytiva, Marlborough, USA). The column was washed with 20 mM imidazole (lysis buffer) and eluted in the same buffer containing 250 mM imidazole. Immediately after purification, the His-tag was cleaved at 4^°^C for 20 h using a recombinant His-tagged Tobacco Etch Virus (TEV) protease. A second IMAC step (IMAC-II) was applied to remove the protease, the uncut protein and the released affinity tag.

The purified proteins were then dialyzed against a 15 mM HEPES pH 7.0, 100 mM NaCl, and 2 mM TCEP buffer. The protein concentration was measured by UV absorption spectroscopy (280 nm) using a NanoDrop 1000 spectrophotometer (Thermo Scientific, Waltham, MA, USA). The purified proteins were concentrated using Amicon Ultra filters (Millipore, Bedford, MA, USA) to 48 mg/ml (L1 MBL) and 50 mg/ml (DBL). Individual aliquots of purified proteins were flash cooled in liquid nitrogen and stored at -80 ^°^C until use.

### 3.2. L1 MBL batch crystallization

We optimized batch crystallization under conditions established previously ^30^. The L1 protein used for batch crystallization was exchanged to a buffer containing 15 mM Tris, 0.1 M KCl, 1.5 mM TCEP, 5 mM ZnCl_2_ pH 7.0 and 200 μl of L1 was added to 200 μl of crystallization buffer: 0.15 M sodium malonate pH 8.0, 20 % (w/v) PEG3350. Crystallizations were set-up in 500 μl polypropylene tubes, which were stored in horizontal position at 289K. The crystals were harvested for SSX approximately after two weeks from setting up crystallization. One hour before SSX data collection crystals were centrifuged at 100 rpm at room-temperature, excess of buffer was removed until 30 μl were left in a tube. The crystals were resuspended in remaining volume using a pipette tip with a cut end to prevent mechanical damage of crystals. Finally, 15 μl of crystals suspension were pipetted evenly on a nylon mesh that was docked in a partially assembled *ALEX* chip. Subsequently, the crystals were covered with a piece of mylar, and the chip was sealed by the top part of *ALEX* holder.

### 3.2. DBL crystallization in microfluidic droplets

The DBL protein was crystallized in aqueous droplets in fluorinated oil using a microfluidic setup. The microfluidic droplet generator chips were designed with the AutoCAD software (Autodesk) and were fabricated via soft lithography at the Center for Nanomaterials (Argonne, IL, USA). Briefly, SU-8 2025 photoresist (MicroChem, Westborough, MA, USA) was used to make molds on a 4-inch silicon wafer. The two-part silicone elastomer (SYLGARD 184, Thermo Fisher Scientific, Waltham, MA, USA), the silane precursor and curing agent were mixed 10:1, degassed, poured on silicon wafer mold, degassed and baked at 65 ^°^C for 4 h. The polydimethylsiloxane (PDMS) elastomer was punched and bonded to a 48 × 65 mm, thickness No. 1 cover glass (Ted Pella Inc., Redding, CA, USA). The droplet generator devices were treated with Aquapel. Two 500 μl glass syringes (Hamilton, Reno, NV, USA) partially filled with droplet generation oil for probes (Bio-Rad, Hercules, CA, USA) and attached to 1/16” OD, 0.02” ID fluorinated ethylene propylene (FEP) tubes (IDEX Corporation, Lake Forest, IL, USA). The oil was pushed from the syringe into each tube and placed in a PHD Ultra Series pump (Harvard Apparatus MA, USA). The screen solutions were prepared from stock reagents supplied by Rigaku (Rigaku Americas Corporation, The Woodlands, TX, USA) and were filtered with a 0.2 μm nylon filter prior use. Small aliquots (10 μl) of the 50 mg/ml protein stock solution (in 20 mM HEPES–NaOH pH 8.0, 250 mM NaCl, 2 mM DTT) and crystallization screen (0.1 M Bis-Tris pH 9.0, 8 % (w/v) PEG 20,000) were drawn into the tubes in parallel at 5 μl min^-1^ flow rate. 23-gauge pins (New England Small Tube Corporation, NE, USA) were used to connect the FEP tubes to the inlets on a PDMS microfluidic chip. A second PHD Ultra Series pump was equipped with a 1 ml glass syringe and filled with Bio-Rad droplet generation oil for probes. The oil, protein and crystallization solution flow rates were set at 10 μl/min and 2 μl/min, respectively. The droplets were collected in a 0.6 ml test tube and incubated at 16 ^°^C. Droplet sizes were controlled by the adjusting relative flow rates and chips with different geometries. Crystals of 10-100 μm were observed in droplets (50-100 μm diameter) in as little as 2 h after droplet generation. The protein concentration was optimized for crystallization in droplets (50 mg/ml) and was higher than what was used in plates (40 mg/ml). The droplet slurry was kept at 16 ^°^C for 4 days until data collection when 30 μl aliquot of crystals suspension was spread onto the *ALEX* holder as described above for the L1 protein (Supplementary figure 2).

### 3.3 Data collection

#### 3.3.1 Serial synchrotron crystallography data collection

For the data collection from crystals of L1 MBL and DBL, we used the *ALEX* holder with window size 12.4 × 9.4 mm. X-ray beam was an unattenuated and collimated to 50 × 50 μm with a 50 μm motor step size and an exposure area of 175 × 220 steps in x and y respectively. The crystal crystal-to-detector distance was 350 mm. The total number of images from the scan was 38,500 for L1 MBL, with the Pilatus3 × 6M detector collecting images with 50 ms exposure time at each scanning point. The total number of images can vary slightly between samples due to stoppages and different starting points. For the DBL from *C. pinensis*, data from only one *ALEX* chip was required to get a structure. For this protein we chose a 75 × 75 μm beam, 50 μm overlapping step, 40 ms exposure time and a 175 × 208 grid resulting in 36,400 images.

Data collection was performed by using an easy-to-use MEDM/python graphical user interface (GUI) developed at the SBC called *Cris*.*py*. The software takes user values and translates them to the motion control program inside the PMAC motor controller. *Cris*.*py* creates two JSON files containing beamline metadata and the user-supplied information about the sample. Therefore, *Cris*.*py* generates all information needed for downstream analysis such as unit cell (UC) dimensions, crystal symmetry, protein name, PDB accession code structure used for molecular replacement, experimental intent, and principal investigator. Data collection statistics are summarized in Table 1.

**Table 1.** Experimental set up and SSX data collection statistics.

| **Proteins** | **DBL**<br>**from *C. pinensis*** | **L1 MBL**<br>**from *S. maltophilia*** |
| --- | --- | --- |
| Beamline | 19-ID, APS | 19-ID, APS |
| Wavelength (Å) | 0.9792 | 0.9792 |
| Temperature (K) | 295 | 295 |
| Detector | PILATUS3 X 6M | PILATUS3 X 6M |
| Beam size (µm) | 75 × 75 | 50 × 50 |
| Step size (µm) | 50 | 50 |
| Flux (ph/s) | 3 × 10<sup>12</sup> | 3 × 10<sup>12</sup> |
| Exposure (ms) | 40 | 50 |
| Crystal – detector distance (mm) | 325 | 350 |
| Number of chips | 1 | 1 |
| Number of collected images | 36,400 | 38,500 |
| Number of indexed images | 8,424 | 7,186 |
| Indexing hit rate (%) | 23.1% | 18.6% |
| Diffraction images rejected with too low resolution | 565 (2.5 Å)<sup>a</sup> | 788 (2.8 Å) |
| Diffraction images with too few reflections | 82 (25) | 218 (75) |
| Diffraction images greater deviation from the crystal UC vol. | 404 (3.5%) | 212 (3.5%) |
| Number of images after rejection of outliers | 7,261 | 6,183 |
| Images with indexed single lattice | 5,758 | 5,814 |
| Images with indexed two lattices | 1293 | 349 |
| Images with indexed three lattices | 210 | 20 |
| **Lattices merged for structure determination** | 7,018 | 5,437 |
| **Total time of data collection (min)** | 80 | 87 |
| **Average diffraction weighted dose (kGy)** | 11.8 | 19.4 |

<sup>a</sup> Values in parentheses correspond to cut-off values

#### 3.3.2 Kanzus data analysis pipeline

We developed a data analysis pipeline, Kanzus, to bridge the beamline with high-performance computing (HPC) and storage capabilities provided by Argonne’s Leadership Computing Facility (ALCF) (Fig. 3). This pipeline manages the transfer of data, orchestration of analysis and visualization tools, and the publication of results to a user-friendly data portal. Prior to the experiment, we initiate the APS’s Data Management (DM) ^33^ system to monitor the experimental data directory and replicate data from the beamline’s local directory to a Globus accessible storage resource. This storage is connected to the ALCF by two 100 Gigabit fiber connections to accommodate rapid transfer of data between the facilities. At the ALCF we employ the Theta supercomputer to accelerate analysis and provide rapid feedback and results during the experiment. Theta is an 11.7 petaflop machine based on Intel Xeon Phi “Knights Landing” (KNL), 64-core processors. Each of the 4,392 nodes contains 16 GB MCDRAM and 192 GB of DDR4 RAM and leverages a high-speed InfiniBand interconnect.

We use the DM system to reliably monitor local storage, replicate data, and trigger the Kanzus pipeline. As DM detects the creation of new files it automatically copies them to an APS-hosted storage device, where they are made available through Globus ^34^. The DM system is configured to initiate the Kanzus pipeline to act on batches of 256 images. The Kanzus pipeline is implemented as a Globus Flow ^35^. Globus Flows is a platform-as-a-service to create, share, and run distributed data management and analysis pipelines. The Kanzus pipeline leverages the Globus Transfer ^34^ service to move data and the funcX ^36^ function as a service platform to reliably and securely perform remote execution. FuncX builds upon the Parsl ^37^ parallel scripting library to interface with heterogeneous computing resources, allowing funcX to acquire compute nodes on the Theta supercomputer. We have implemented funcX functions for the DIALS ^38^ and PRIME ^39^ suites of crystallography tools with custom visualization Python scripts *Rejectoplot* and *Primalysis* to enable their use within the Kanzus pipeline.

In the case of the DBL datasets, the Kanzus pipeline performed stills processing to identify crystal lattices and integrate diffraction intensity, visualized intermediate hit-rates and metrics during the chip’s processing, and submitted results to the beamline’s data portal. Two metadata JSON files created for each collection run are placed into the directory prior to data landing there and are therefore part of the first 256 files to be transferred. These metadata files are used as input to the analysis and are ingested as metadata to the portal. Since these two datasets were collected, we have extended the Kanzus pipeline to incorporate beam X-Y search capabilities to automatically search for optimal XY coordinates for the incident beam to use during analysis, and to refine the structure using the PRIME tool ^39^. Later, we aim to incorporate pre-processing and triaging steps to remove unnecessary data and accelerate the time to solution.

#### 3.3.3 Structure refinement

Integrated (*int*) files from the DIALS *stills_process* were returned to the local computers for merging and scaling steps prior to the Kanzus pipeline upgrade a few months later. The data reported here were merged and scaled by consecutive rounds of PRIME and two new pieces of software, *Rejectoplot* and *Primalysis*. PRIME is a bulk refinement that scales all the integrated and indexed reflections. *Rejectoplot* and *Primalysis* were developed to interrogate the outputs of PRIME, skimming outliers (*Rejectoplot*) and making estimations for resolution cut-off (*Rejectoplot*). These two scripts are integrated into the Kanzus pipeline as an autonomous decision point. All processed SSX data in intensities (I) were converted to structure factor amplitudes (F) in the Ctruncate program ^40,41^ from the CCP4 package ^42^ before being used for structure determination.

Both structures were solved by molecular replacement using Molrep ^43^ with one subunit of the reported L1 structure (PDB id 6UA1) for L1 and the previously refined structure of the same protein for the Class D *β* -lactamase, respectively, as search models. These initial models were then refined by 5 cycles of rigid-body refinement followed by additional 10 cycles of restrained refinement using Refmac ^44^. The structures were further refined iteratively until converged with reasonable stereochemistry by using Phenix ^45^ for computational refinement and Coot ^46^ for manual adjustment. The progress of the refinement was monitored by inspecting R and R_free_ values, Ramachandran plots and MOLPROBITY ^47^ throughout the process. The final structures of L1 (24-287 with two zinc ions) and DBL (23-267) were then validated with RCSB validation service. The atomic coordinates and structure factors have been deposited in the Protein Data Bank under accession codes 7L52 and 7K3M for L1 and Class D *β* -lactamase, respectively. The final refinement statistics are given in Table 2.

**Table 2.** Data collection and structure refinement statistics.

|  | DBL<br>from *C. pinensis* | L1 MBL<br>from *S. maltophilia* |
| --- | --- | --- |
| PDB ID | 7K3M | 7L52 |
| Space group | *P2<sub>1</sub>2<sub>1</sub>2<sub>1</sub>* | *P6<sub>4</sub>22* |
| a, b, c (Å) | 49.41, 69.35, 70.04 | 105.67, 105.67, 99.33 |
| Resolution range (Å)<sup>a</sup> | 49.28 – 1.80 (1.83 – 1.80) | 46.65 – 1.85 (1.88 – 1.85) |
| No. of reflections | 23,276 (1,134) | 28,773 (1,398) |
| Completeness (%) | 99.99 (99.82) | 99.99 (100.00) |
| Data redundancy | 79.11 (36.99) | 57.37 (29.22) |
| R<sub>split</sub> (%)<sup>b</sup> | 15.19 (40.49) | 19.75 (36.24) |
| CC<sub>1/2</sub><sup>c</sup> | 0.973 (0.732) | 0.942 (0.818) |
| $\langle I/\sigma(I) \rangle$ | 2.81 (0.76) | 4.08 (1.15) |
| Wilson *B* factor | 11.5 | 6.93 |
| **Structure Determination** |  |  |
| MR initial model (PDB ID) | complex with avibactam | 6UA1 |
| **Refinement** |  |  |
| Resolution range (Å) | 49.28 – 1.80 (1.88 – 1.80) | 46.65 – 1.85 (1.92 – 1.85) |
| Completeness (%) | 100.0 (95.0) | 100.0 (100.0) |
| No. of reflections | 22,940 (2667) | 28,496 (2793) |
| R<sub>work</sub>/R<sub>free</sub><sup>d</sup> (%) | 0.178/0.224(21.72 /22.53) | 17.52 /21.67(22.65 /25.99) |
| Protein chains/atoms | 2,029 | 1,998 |
| Ligand/Solvent atoms | 131 | 179 |
| Mean temperature factor (Å<sup>2</sup>) | 15.62 | 13.42 |
| **Coordinate Deviations** |  |  |
| R.m.s.d. bonds (Å) | 0.003 | 0.006 |
| R.m.s.d. angles (°) | 0.580 | 0.761 |
| **Ramachandran plot**<sup>e</sup> |  |  |
| Favored (%) | 96.0 | 96.0 |
| Allowed (%) | 4.0 | 4.0 |
| Outside allowed (%) | 0 | 0 |

<sup>a</sup> Values in parentheses correspond to the highest resolution shell.<sup>b</sup> R<sub>split</sub> as defined by White <sup>48</sup>.<sup>c</sup> As defined by Karplus and Diederichs <sup>49</sup>.<sup>d</sup> $R = \Sigma |F_o| - |F_c| / \Sigma |F_o|$ for all reflections, where *F<sub>o</sub>* and *F<sub>c</sub>* are observed and calculated structure factors, respectively. R<sub>free</sub> is calculated analogously for the test reflections, randomly selected, and excluded from the refinement.<sup>e</sup> As defined by Molprobity <sup>47</sup>

#### 3.3.4 Updates to the SSX set-up

The original design of the SSX system is undergoing continuous improvements. Since the initial implementation and proof-of-concept experiments, a number of features have been updated. As previously mentioned in section 2.1, since the data mentioned here was collected the serial motors have been permanently integrated onto the goniometer. This reduced the transition time from single-crystal experiments to serial experiments from 30 minutes both ways, to less than 5 minutes. This reduction in transition time now makes performing multiple experiment types in the same scheduled beamtime much more feasible as there is no longer a significant downtime. The serial motors are installed in a modular fashion which can be removed and reattached with high precision, allowing for adding-on of an already designed plates canner system or other future equipment. This design also had the added benefit of improving the center-of-rotation alignment due to being lighter weight with less of a lever arm.

The second major addition to the beamline system was a compact refractive lens system. This system can focus the beam to approximately 15 × 5 μm and can be aligned and ready for use in under half an hour. This system greatly increases the flux density in the focal spot and can be used for both serial and single crystal crystallography. With the smaller beam size and higher flux density, more images can be collected from an *ALEX* chip with the same quantity of sample.

## 4. Results

### 4.1. Development of the SSX capability at SBC

SSX has become a valuable structural biology approach to study chemical transformations and dynamic processes in protein crystals. Sample limitations and crystal delivery to the x-ray beam represent important challenges for this approach to become more accessible to the broader scientific community. In response to the growing need for SSX and to address existing bottlenecks we have developed a new, fixed-target method implemented at the Structural Biology Center. Initial studies were conducted on crystals obtained in microfluidic droplets (manuscript in review). These successful experiments demonstrated that our SSX method is robust, performs well, and is ready to be rolled out to general users.

The sample environment is provided by the ALEX holder, a small area mylar-mesh-mylar sandwich designed to enable collection of a complete diffraction data set from a single chip. The holder immobilizes and seals a near-monolayer of crystals on a nylon mesh. The developed method is simple and low-cost, accepts crystals of various sizes, and can be applied to both static and time-resolved SSX. The holders are attached onto the end-station goniostat using a magnetic mount in the same fashion as a standard single crystal crystallography pin, and data collection is controlled by simple software developed at SBC. The *ALEX* accepts small volume of microcrystal suspension which can be shipped in vials and loaded at the synchrotron and set up for data collection on-site by staff or remotely by the user. The *ALEX* allows data collection from tens of thousands of crystals while maintaining a biologically relevant temperature. In many cases data collected from a single chip are sufficient for structure determination. It provides reliable, user-friendly access to serial crystallography methods. The *ALEX* is flexible enough to transition quickly between different types of experiments.

### 4.2 Quality of SSX diffraction

Due to the physical geometry of the 19-ID beamline and the nature of the *ALEX* holder, the current method of fixed-target serial synchrotron crystallography is not optimized for low-background data collection. Much of the background scattering occurs from the beam passing through the relatively dense nylon mesh used to immobilize the crystals, of which nylon fiber diffraction can be seen in varying amounts in every diffraction image (Fig. 3). Other sources of background scattering come from the beamline itself due to its orientation and endstation components, meaning that these sources could be reduced with an improved set-up and component changes.

### 4.3 Data processing and structure solution

The L1 MBL had an effective hit rate of 18.6 % and 7186 indexed images were recovered. Outlier rejection was performed by analyzing the output from the post-refinement software PRIME, where images were rejected if they failed any of these three criteria: i) Less than 75 diffraction spots per image, ii) Resolution worse than 2.8 Å and iii) Unit cell dimensions which were greater or less than 3.5 % of the average (see supplementary graph 1 (*rejectoplot*)). The resulting 6183 images were then subjected to another round of post-refinement with PRIME using space group P6_4_22. The resolution cut-off at 1.85 Å was used following the three main metrics, I**2 rapidly increasing above 2, CC_1/2_ falling below 0.5, and completeness dropping to 99.9 % (see supplementary graph 2 (*primalysis*)).

The DBL had a higher effective hit rate of 23.1% with total of 8424 diffraction images prior to outlier rejection. Outlier rejection parameters were the same except for images being rejected if they had less than 25 diffraction spots. The change from 75 to 25 was due to the unit cell being significantly smaller, resulting in fewer spots per image. A total of 1163 images were rejected, resulting in 7261 moving into the final merging step. The DBL had a space group of P2_1_2_1_2_1_. Resolution cut off metrics were the same as with the L1 MBL giving a final resolution of 1.80 Å.

The two structures were determined by molecular replacement and refined and validated as described in Methods. DBL from *C. pinensis* is built by a large anti-parallel β-sheet decorated by 9 α-helices (Fig. 5A), while L1 from *S. maltophilia* possesses canonical fold for MBLs with hydrophobic core made by two β-sheet inserted between α-helices (Fig. 5B). Both structures are first room temperature models of these proteins and demonstrate great agreement with their cryogenic equivalents, including organization of their active sites. The two SSX structures superpose with their cryo counterparts with overall root mean square deviation (RMSD) of Cα atoms = 0.28 Å over 264 residues for L1 MBL and 0.292 Å over 245 residues for DBL. The structural details will be described elsewhere.

**Figure 4.**
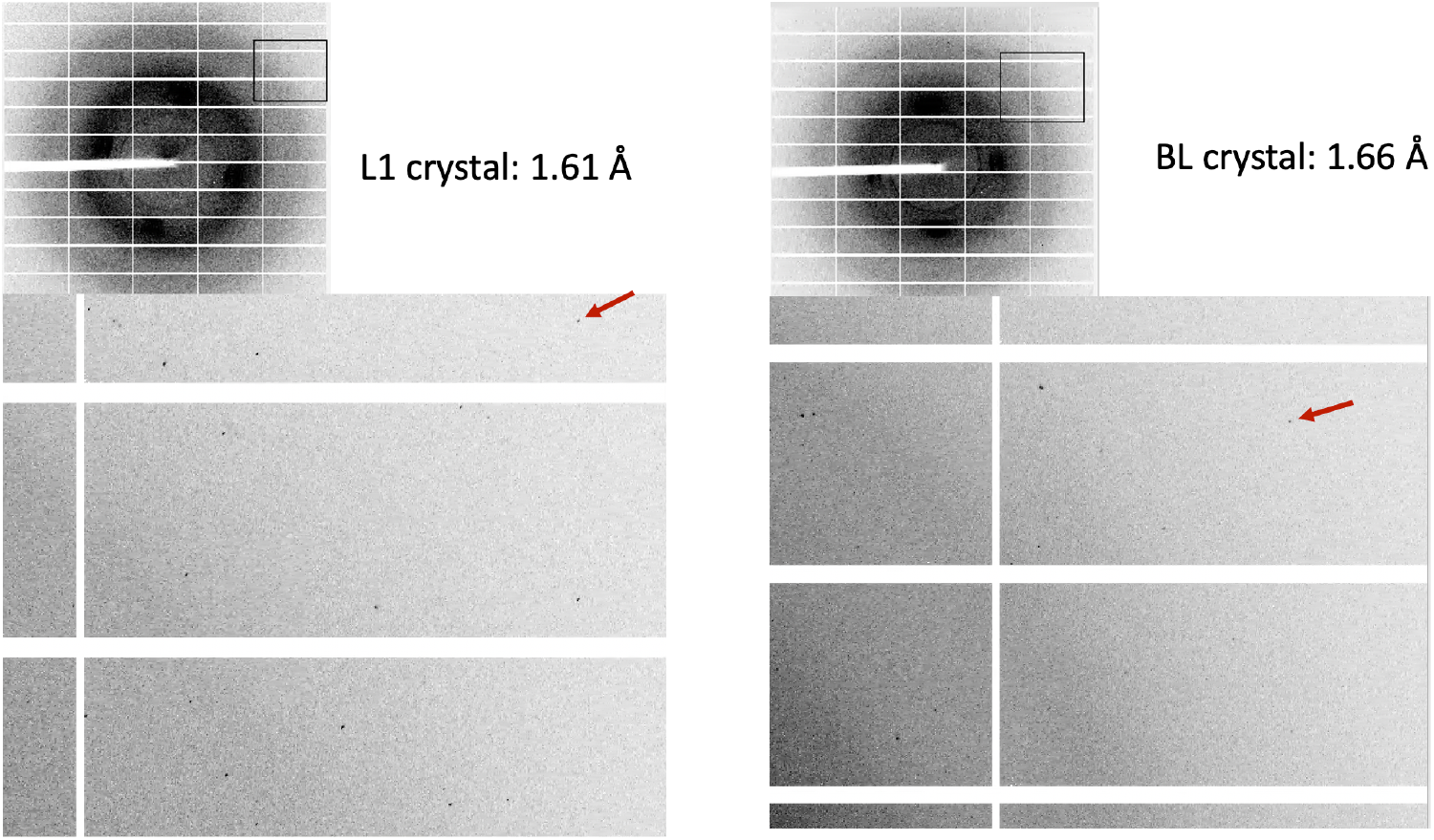
Diffraction patterns obtained from crystals of L1 (A) and DBL (B). The arrows point to reflection at indicated resolution.

**Figure 5.**
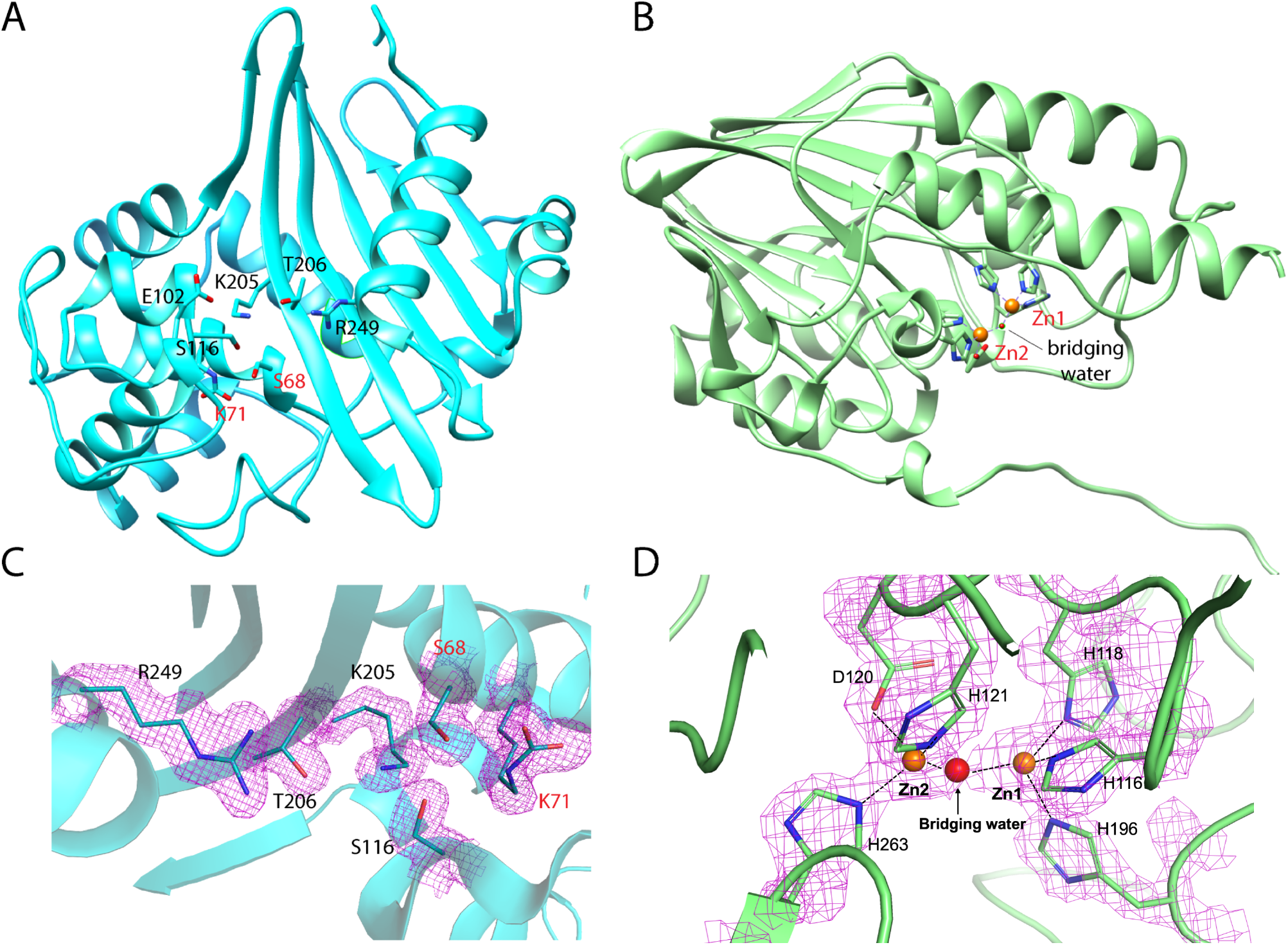
Room-temperature SSX crystal structures of class B and D β-lactamases. (A) Crystal structure of DBL from *C. pinensis* determined at 1.80 Å. (B) Crystal structure of class L1 MBL from *S. maltophilia* determined at 1.85 Å. 2mFo-DFc electron density maps calculated for active sites of *C. pinensis* DBL (C), and L1 MBL (D). Difference electron density maps contoured at 1.2 σ level.

Quality of the SSX room-temperature data is illustrated by the 2mFo-DFc maps contoured around residues in the active site of DBL from *C. pinensis* and L1 MBL from *S. maltophilia* (Figs. 5C, D). Visual inspection of the calculated 2mFo-DFc maps around catalytic residues shows that maps are comparable or better than cryo-crystallography structures solved at similar resolution. The system for SSX data collection is ready for structural dynamic studies of β-lactamases and other enzymes.

## 5. Conclusions

Serial crystallography has become available at XFELs and now also at light sources around the world. It provides an exciting opportunity to study enzyme structures under physiological temperatures and lower radiation dose and investigate structural dynamics using crystallography. We responded to the increasing demand for SSX, and we have developed a simple and accessible fixed-target serial crystallography approach. The key elements of this system include: (i) the reusable, 3D-printed fixed-target *ALEX* sample holder for the immobilization of the crystal suspension and (ii) a robust computational pipeline for data transfer and processing utilizing the high-performance computing resources to generate on-the-fly feedback about data quality. The applicability of the design has been demonstrated on two model proteins, which yielded first room-temperature structures for these macromolecules with quality comparable to their cryogenic counterparts.

This SSX data collection system was implemented at the SBC 19-ID beamline at the APS. To complement the data collection, we have established the Kanzus software data processing pipeline linked to the high-performance computing resources at the ALCF. The pipeline moves the data to Globus accessible storage resource and executes analysis of diffraction images providing feedback to data collection via user-friendly data portal.

The SSX implemented at SBC was designed with general users in mind. The goal was to make SSX easy to access, reduce the cost of use, and be simple enough as to not create a technological barrier preventing its wider adoption. The beamline systems and the *ALEX* holder accomplish these goals by having a specialized holder that can be 3D printed in under an hour (protocol for 3D printing is available on request) for less than $20 in material cost, and a beamline that is flexible enough to transition quickly between different types of experiments. Our system can be applied to both static and time-resolved studies.

To provide access to general users, crystals suspension can be shipped in vials which can then be loaded on-site by SBC staff with data collection being performed by staff or remotely by a user. Other options can include users traveling on-site and performing these actions themselves with staff oversight or having fully loaded and assembled *ALEX* holders shipped in for remote data collection. The latter option would require creation of a shipping box which would keep the samples viable during transport. The SBC is now accepting general user proposals for beamtime in a collaborative mode.

## Acknowledgements

We would like to thank Paula Bulaon for help with preparing this manuscript. Funding for this project was provided in part by federal funds from the National Institute of Allergy and Infectious Diseases, National Institutes of Health, Department of Health and Human Services, under Contracts No. HHSN272201200026C and HHSN272201700060C. The use of SBC beamlines at the Advanced Photon Source is supported by the U.S. Department of Energy (DOE) Office of Science and operated for the DOE Office of Science by Argonne National Laboratory under Contract No. DE-AC02-06CH11357.

## Supplementary materials

### Protocol for performing fixed target SSX experiment at 19-ID

This section provides a brief overview of necessary steps required in a live experiment. While many of the specifics of each step are provided in the above sections of this paper, below is a summary of the steps in order with a brief description:

1. Perform typical beam setup including tune, guard slit scans, and beam alignment to center-of-rotation crosshairs.
  a. If CRLs are being used, beamline monochromator sagittal focus, vertical focusing mirror, downstream supports, guardslit supports, and goniometer supports must all be moved to pre-identified values (10-20 minutes)
  b. CRL lens insertion/extraction motor must then be moved into place, and monochromator tuning and guardslit tuning must be performed
  c. After a and b are completed, move beam to center-of-rotation cross hairs for use.
2. Transition beamline to Serial Mode:
  a. Disable auto-mounting robot
  b. Retract cold stream
  c. Move Gonio X prime motor as necessary
3. Ensure Data Management client is running to transfer data to supercomputing for processing
4. Load crystals into ALEX holder
5. Mount ALEX holder and align to bottom left corner
  a. In most instances, running a small test with burn paper is performed to ensure system is working properly.
6. In the *Crys*.*py* GUI, load all fields including crystals space group, PDB, sample name, etc.
7. Calculate and enter step size, beam size, exposure time, and detector distance into GUI
8. Press *Arm* button in GUI, which loads all necessary parameters to beamline components for collection.
9. Press *Collect* button after Arm procedure completes
10. Monitor first rows to ensure system is performing nominally
11. Repeat steps 4-10 until experiment is complete
12. After last serial sample is complete reset beamline by:
  a. Moving Gonio X prime motor to previous position
  b. Insert cold stream to previous position (0 mm)
  c. Reenable auto-mounting robot

**Supplementary Figure S1.**
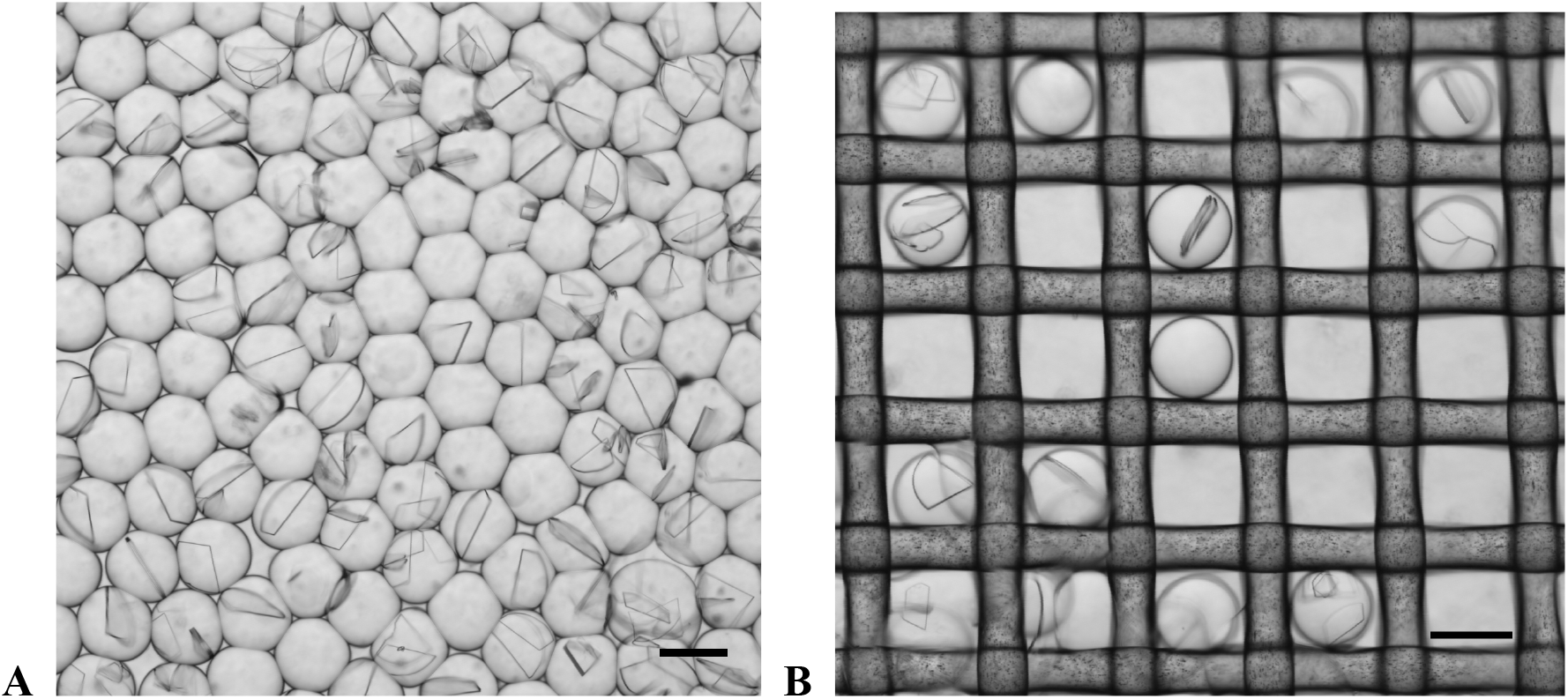
Droplet generator chips were used to generate 45-55 μm aqueous droplets in fluorinated oil. Varying protein concentration and flow ratios (protein, precipitant and oil) was used to optimize droplet size, occupancy and number of crystals in the droplet (A). 30 μl of droplet slurry was pipetted evenly on a nylon mesh in the partially assembled ALEX chip(B). Scale bar: 50 μm.

## References

1. Chapman, H. N. X-Ray Free-Electron Lasers for the Structure and Dynamics of Macromolecules. Annual Review of Biochemistry 88, 35–58, doi:10.1146/annurev-biochem-013118-110744 (2019).

2. Coe, J. & Fromme, P. Serial femtosecond crystallography opens new avenues for Structural Biology. Protein & Peptide Letters 23, 255–272, doi:http://dx.doi.org/10.2174/0929866523666160120152937 (2016).

3. Schlichting, I. Serial femtosecond crystallography: the first five years. IUCrJ 2, 246–255, doi:doi:10.1107/S205225251402702X (2015).

4. Pei, K. L., Sooriyaarachchi, M., Sherrell, D. A., George, G. N. & Gailer, J. Probing the coordination behavior of Hg2+, CH3Hg+, and Cd2+ towards mixtures of two biological thiols by HPLC-ICP-AES. Journal of Inorganic Biochemistry 105, 375–381, doi:https://doi.org/10.1016/j.jinorgbio.2010.11.019 (2011).

5. Wiedorn, M. O. et al. Megahertz serial crystallography. Nature Communications 9, 4025, doi:10.1038/s41467-018-06156-7 (2018).

6. Martiel, I., Müller-Werkmeister, H. M. & Cohen, A. E. Strategies for sample delivery for femtosecond crystallography. Acta Crystallogr D Struct Biol 75, 160–177, doi:10.1107/S2059798318017953 (2019).

7. de la Mora, E. et al. Radiation damage and dose limits in serial synchrotron crystallography at cryo- and room temperatures. Proceedings of the National Academy of Sciences 117, 4142, doi:10.1073/pnas.1821522117 (2020).

8. Mehrabi, P. et al. The HARE chip for efficient time-resolved serial synchrotron crystallography. Journal of Synchrotron Radiation 27, 360–370, doi:doi:10.1107/S1600577520000685 (2020).

9. Pearson, A. R. & Mehrabi, P. Serial synchrotron crystallography for time-resolved structural biology. Current Opinion in Structural Biology 65, 168–174, doi:https://doi.org/10.1016/j.sbi.2020.06.019 (2020).

10. Weinert, T. et al. Proton uptake mechanism in bacteriorhodopsin captured by serial synchrotron crystallography. Science 365, 61–65, doi:10.1126/science.aaw8634 (2019).

11. Wilamowski, M. et al. 2′-O methylation of RNA cap in SARS-CoV-2 captured by serial crystallography. Proceedings of the National Academy of Sciences 118, e2100170118, doi:10.1073/pnas.2100170118 (2021).

12. Mishin, A. et al. An outlook on using serial femtosecond crystallography in drug discovery. Expert Opinion on Drug Discovery 14, 933–945, doi:10.1080/17460441.2019.1626822 (2019).

13. Orville, A. M. Recent results in time resolved serial femtosecond crystallography at XFELs. Current Opinion in Structural Biology 65, 193–208, doi:https://doi.org/10.1016/j.sbi.2020.08.011 (2020).

14. Doak, R. B. et al. Crystallography on a chip - without the chip: sheet-on-sheet sandwich. Acta Crystallogr D Struct Biol 74, 1000–1007, doi:10.1107/S2059798318011634 (2018).

15. Hunter, M. S. et al. X-ray Diffraction from Membrane Protein Nanocrystals. Biophysical Journal 100, 198–206, doi:https://doi.org/10.1016/j.bpj.2010.10.049 (2011).

16. Lee, D. et al. Nylon mesh-based sample holder for fixed-target serial femtosecond crystallography. Scientific Reports 9, 6971, doi:10.1038/s41598-019-43485-z (2019).

17. Lyubimov, A. Y. et al. Capture and X-ray diffraction studies of protein microcrystals in a microfluidic trap array. Acta Crystallogr D Biol Crystallogr 71, 928–940, doi:10.1107/S1399004715002308 (2015).

18. Oberthuer, D. et al. Double-flow focused liquid injector for efficient serial femtosecond crystallography. Scientific Reports 7, 44628, doi:10.1038/srep44628 (2017).

19. Roedig, P. et al. A micro-patterned silicon chip as sample holder for macromolecular crystallography experiments with minimal background scattering. Scientific Reports 5, 10451, doi:10.1038/srep10451 (2015).

20. Echelmeier, A. et al. 3D printed droplet generation devices for serial femtosecond crystallography enabled by surface coating. J Appl Crystallogr 52, 997–1008, doi:doi:10.1107/S1600576719010343 (2019).

21. Schulz, J. et al. A versatile liquid-jet setup for the European XFEL. Journal of synchrotron radiation 26, 339–345, doi:10.1107/S1600577519000894 (2019).

22. Fromme, R. et al. Serial femtosecond crystallography of soluble proteins in lipidic cubic phase. IUCrJ 2, 545–551, doi:10.1107/S2052252515013160 (2015).

23. Beyerlein, K. R. et al. Mix-and-diffuse serial synchrotron crystallography. IUCrJ 4, 769–777, doi:10.1107/S2052252517013124 (2017).

24. Weinert, T. et al. Serial millisecond crystallography for routine room-temperature structure determination at synchrotrons. Nature Communications 8, 542, doi:10.1038/s41467-017-00630-4 (2017).

25. Owen, R. L. et al. Low-dose fixed-target serial synchrotron crystallography. Acta Crystallogr D Struct Biol 73, 373–378, doi:10.1107/S2059798317002996 (2017).

26. Rosenbaum, G. et al. The Structural Biology Center 19ID undulator beamline: facility specifications and protein crystallographic results. Journal of synchrotron radiation 13, 30–45, doi:10.1107/S0909049505036721 (2006).

27. Shu, D. et al. Mechanical Design of a New Precision Alignment Apparatus for Compact X-ray Compound Refractive Lens Manipulator \WEOPMA04, doi:10.18429/JACoW-MEDSI2018-WEOPMA04 (2018).

28. Ebrahim, A. et al. Dose-resolved serial synchrotron and XFEL structures of radiation-sensitive metalloproteins. IUCrJ 6, 543–551, doi:10.1107/S2052252519003956 (2019).

29. Ebrahim, A. et al. Resolving polymorphs and radiation-driven effects in microcrystals using fixed-target serial synchrotron crystallography. Acta Crystallographica Section D 75, 151–159, doi:doi:10.1107/S2059798318010240 (2019).

30. Kim, Y. et al. Structural and biochemical analysis of the metallo-β-lactamase L1 from emerging pathogen Stenotrophomonas maltophilia revealed the subtle but distinct di-metal scaffold for catalytic activity. Protein Sci 29, 723–743, doi:10.1002/pro.3804 (2020).

31. Donnelly, M. I. et al. An expression vector tailored for large-scale, high-throughput purification of recombinant proteins. Protein Expression and Purification 47, 446–454, doi:https://doi.org/10.1016/j.pep.2005.12.011 (2006).

32. Kim, Y. et al. High-throughput protein purification and quality assessment for crystallization. Methods 55, 12–28, doi:http://dx.doi.org/10.1016/j.ymeth.2011.07.010 (2011).

33. Veseli, S., Schwarz, N. & Schmitz, C. APS Data Management System. Journal of Synchrotron Radiation 25, 1574–1580, doi:doi:10.1107/S1600577518010056 (2018).

34. Chard, K., Tuecke, S. & Foster, I. Efficient and Secure Transfer, Synchronization, and Sharing of Big Data. IEEE Cloud Computing 1, 46–55, doi:10.1109/MCC.2014.52 (2014).

35. Ananthakrishnan, R. et al. Globus Platform Services for Data Publication. Pearc ‘18, doi:10.1145/3219104.3219127 (2018).

36. Chard, R. et al. FuncX: A Federated Function Serving Fabric for Science. Hpdc ‘20, 65–76, doi:10.1145/3369583.3392683 (2020).

37. Babuji, Y. et al. Parsl: Pervasive Parallel Programming in Python. Hpdc ‘19, 25–36, doi:10.1145/3307681.3325400 (2019).

38. Winter, G. et al. DIALS as a toolkit. Protein Sci 31, 232–250, doi:10.1002/pro.4224 (2022).

39. Uervirojnangkoorn, M. et al. Enabling X-ray free electron laser crystallography for challenging biological systems from a limited number of crystals. eLife 4, e05421, doi:10.7554/eLife.05421 (2015).

40. French, S. & Wilson, K. On the treatment of negative intensity observations. Acta Crystallographica Section A 34, 517–525, doi:doi:10.1107/S0567739478001114 (1978).

41. Padilla, J. E. & Yeates, T. O. A statistic for local intensity differences: robustness to anisotropy and pseudo-centering and utility for detecting twinning. Acta Crystallographica Section D 59, 1124–1130, doi:doi:10.1107/S0907444903007947 (2003).

42. Winn, M. D. et al. Overview of the CCP4 suite and current developments. Acta Crystallogr D Biol Crystallogr 67, 235–242, doi:10.1107/S0907444910045749 (2011).

43. Vagin, A. & Teplyakov, A. Molecular replacement with MOLREP. Acta Crystallographica Section D-Biological Crystallography 66, 22–25, doi:10.1107/S0907444909042589 (2010).

44. Murshudov, G. N. et al. REFMAC5 for the refinement of macromolecular crystal structures. Acta Crystallographica Section D 67, 355–367, doi:doi:10.1107/S0907444911001314 (2011).

45. Adams, P. D. et al. PHENIX: a comprehensive Python-based system for macromolecular structure solution. Acta Crystallogr D Biol Crystallogr 66, 213–221 (2010).

46. Emsley, P. & Cowtan, K. Coot: model-building tools for molecular graphics. Acta Crystallogr D Biol Crystallogr 60, 2126–2132, doi:S0907444904019158 [pii] 10.1107/S0907444904019158 (2004).

47. Davis, I. W., Murray, L. W., Richardson, J. S. & Richardson, D. C. MOLPROBITY: structure validation and all-atom contact analysis for nucleic acids and their complexes. Nucleic Acids Res 32, W615–619, doi:10.1093/nar/gkh39832/suppl_2/W615 [pii] (2004).

48. White, T. A. et al. Crystallographic data processing for free-electron laser sources. Acta Crystallographica Section D 69, 1231–1240, doi:doi:10.1107/S0907444913013620 (2013).

49. Karplus, P. A. & Diederichs, K. Linking Crystallographic Model and Data Quality. Science 336, 1030–1033, doi:10.1126/science.1218231 (2012).

